# *OpenGenomeBrowser*: A versatile, dataset-independent and scalable web platform for genome data management and comparative genomics

**DOI:** 10.1101/2022.07.19.500583

**Authors:** Thomas Roder, Simone Oberhänsli, Noam Shani, Rémy Bruggmann

**Affiliations:** Interfaculty Bioinformatics Unit and Swiss Institute of Bioinformatics, University of Bern, 3012 Bern, Switzerland; Methods Development and Analytics, Agroscope, Schwarzenburgstrasse 161, CH-3003 Bern, Switzerland

**Author notes:** Rémy Bruggmann, Interfaculty Bioinformatics Unit, Baltzerstrasse 6, CH-3012 Bern, Switzerland.

**Keywords:** genome database, genome browser, comparative genomics, open-source, self-hosted

## Abstract

OpenGenomeBrowser is a self-hostable open-source platform that manages access to genomic data and drastically simplifies comparative genomics analyses. It enables users to interactively generate phylogenetic trees, compare gene loci, browse biochemical pathways, perform gene trait matching, create dot plots, execute BLAST searches, and access the data. It features a flexible user management system, and its modular folder structure enables the organization of genomic data and metadata, and to automate analyses. We tested OpenGenomeBrowser with bacterial, archaeal and yeast genomes. The largest instance currently contains over 1,400 bacterial genomes. Source code, documentation, tutorials and a demo server are available at opengenomebrowser.github.io.

## Background

Driven by advances in sequencing technologies, many organizations and research groups have accumulated large amounts of genomic data. As sequencing projects progress, the organization of such genomic datasets becomes increasingly difficult. Systematic ways of storing data and metadata, tracking and denoting changes in assemblies or annotations, and enabling easy access are key challenges. While standardized data formats and free software are widely used in the field to process genomic data, data exploration is often still cumbersome. This is especially true for non-bioinformaticians, although numerous platforms have been developed to simplify data access.

Most of these platforms have different user interfaces and sometimes limited functionality. The reason for this heterogeneity is that most of them have been developed independently, i.e., each one for a specific genomic dataset. Such platforms exist for many well-studied organisms, such as *Pseudomonas spp*. (1), but also for non-model species such as ginseng (2) and cork oak (3). These platforms share a set of core features: access to data, sequence similarity searches (like BLAST (4)), and limited annotation searches. The most advanced of these platforms, such as CoGe (5), MicrobesOnline (6), WormBase (7), Genomicus (8) and ChlamDB (9), include additional functions to answer a wide range of questions.

However, these platforms tend to be tied to the characteristics of a specific dataset and adapting their software to other projects would be extremely difficult. This is surprising given that the underlying data are essentially the same: genome assemblies, genes, proteins, and their annotations. Fortunately, this information is stored in standardized data formats across many fields, which in principle would allow code reuse and collaborative development. Even while some degree of purpose-built software tools may still be necessary for certain projects, independent development comes at a significant initial cost as well as a long-term maintenance cost and a higher risk of becoming outdated.

We addressed these issues by developing OpenGenomeBrowser, a self-hostable, open-source software based on the Python web framework Django (10). OpenGenomeBrowser runs on all modern browser engines (Firefox, Chrome, Safari). It contains more features than most similar platforms, is highly user-friendly and *dataset-independent* – i.e., not bound to any specific genomic dataset.

## Results

To enable automated processing of genomic data, as in OpenGenomeBrowser, it is essential that the data is stored in a systematic fashion. We present our solution to this problem in detail in the section “*folder structure*”. The subsequent section “*OpenGenomeBrowser tools*” describes a set of scripts that simplify the handling of the aforementioned folder structure.

### Folder structure

Every sequencing project faces an important challenge: systematic storage of data and metadata according to the FAIR principles (11). These principles enable reproducibility, automation, data interoperability and sharing. Especially in long-term projects, it is crucial to know when and how the data was generated, and to have a transparent way of handling different genome and annotation versions. Different versions are the result of organism re-sequencing, raw data re-assembly or assembly re-annotation. Importantly, each version of a gene must have a unique identifier, and legacy data should be kept instead of being overwritten.

To address these problems, we developed a modular folder structure (Figure 1A). The *organisms* folder contains a directory for each biological entity, e.g., a bacterial strain. Each of these folders must contain a metadata file, *organism*.*json* (Figure 1A, center), describing the biological entity, and a folder named *genomes*. The *genomes* folder contains one folder for each genome version. One of these genomes must be designated as the *representative* genome of the biological entity in *organism*.*json*. This allows project maintainers to update an assembly transparently, by designating the new version as *representative* without removing the old one.

**Figure 1:**
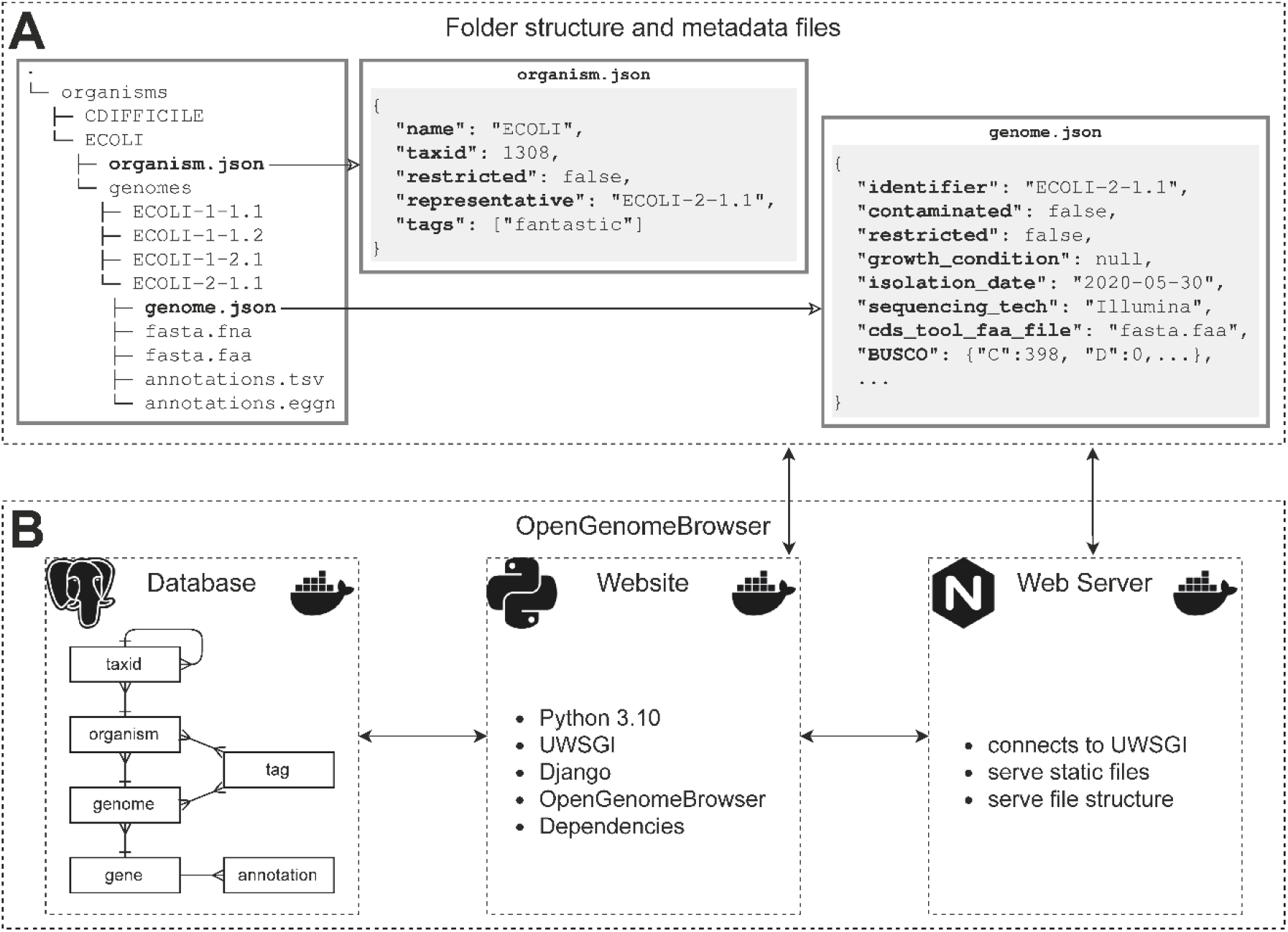
Schematic diagram of OpenGenomeBrowser. (**A**) The user-provided folder structure and metadata files. (**B**) The OpenGenomeBrowser software stack in Docker Compose. It consists of a database container (PostgreSQL), a webserver container (nginx), and a container that executes the OpenGenomeBrowser code.

Each genome folder must contain a metadata file, *genome*.*json* (Figure 1A), and the actual data: an assembly FASTA file, a GenBank file, and a gff3 (general feature format version 3) file. While not strictly required but strongly recommended, annotation files in tab-separated format which map gene identifiers to annotations, may be provided. OpenGenomeBrowser supports several annotation types by default, such as Enzyme Commission numbers, KEGG (12) genes and KEGG reactions, Gene Ontology terms (13,14), and annotations from EggNOG (15). Additional annotation types can be easily configured. Files that map annotations to descriptions (e.g., EC:1.1.1.1 → alcohol dehydrogenase) can be added to a designated folder.

### OpenGenomeBrowser tools

A set of scripts called *OpenGenomeBrowser Tools* simplifies the creation of the previously described folder structure and the incorporation of new genomes. As shown below, a functional folder structure that contains one genome can be set up with only four commands.

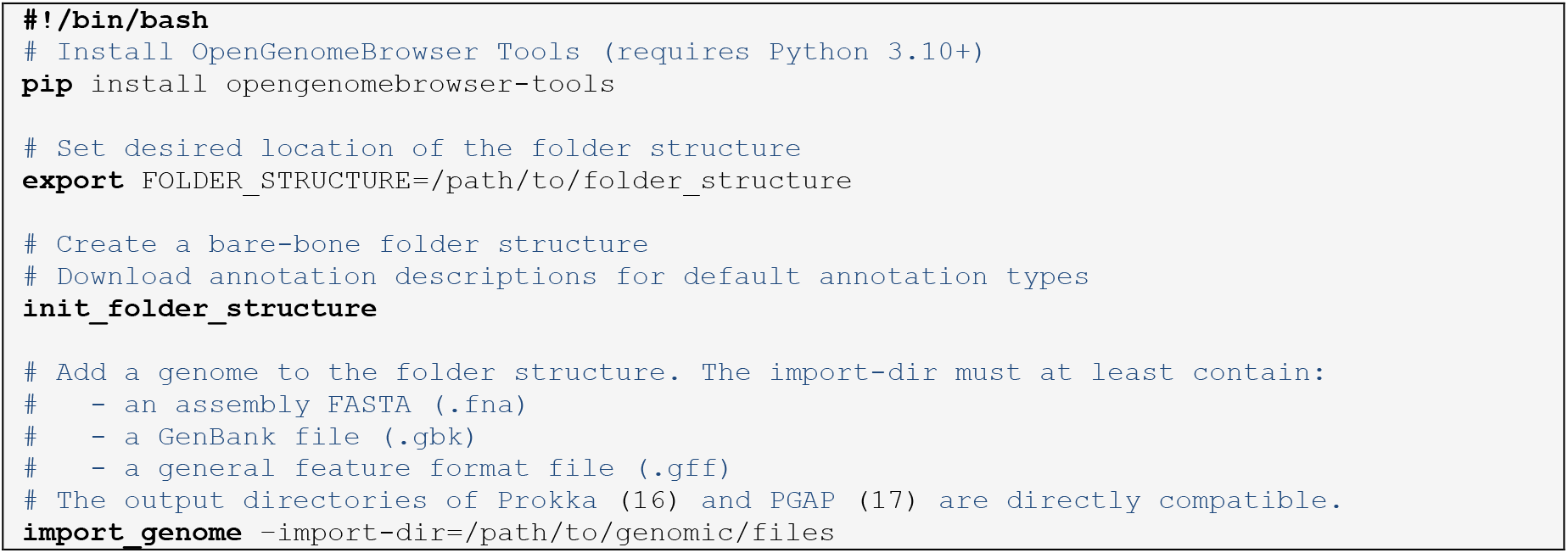

### Software architecture

OpenGenomeBrowser itself is distributed as a Docker container (18). Using Docker Compose, the container is combined with a database and a webserver to create a production-ready software stack (Figure 1B).

#### Features

The following section describes the main features of OpenGenomeBrowser. The reader may try them out at opengenomebrowser.bioinformatics.unibe.ch, where a freely accessible demo server with 24 bacterial genomes is hosted.

### Genomes table

Especially in large sequencing projects, it is vital that the data can be filtered and sorted according to metadata. This is the purpose of the *genomes table view* (Figure 2) which serves as the entry point of OpenGenomeBrowser. By default, only the *representative* genomes are listed and only the name of the organism, the genome identifier, the taxonomic name, and the sequencing technology are shown as columns. Furthermore, there are over forty additional metadata columns available that can be dynamically added to the table. All columns can be used to filter and sort the data, which makes this view the ideal entry point for an analysis.

**Figure 2:**
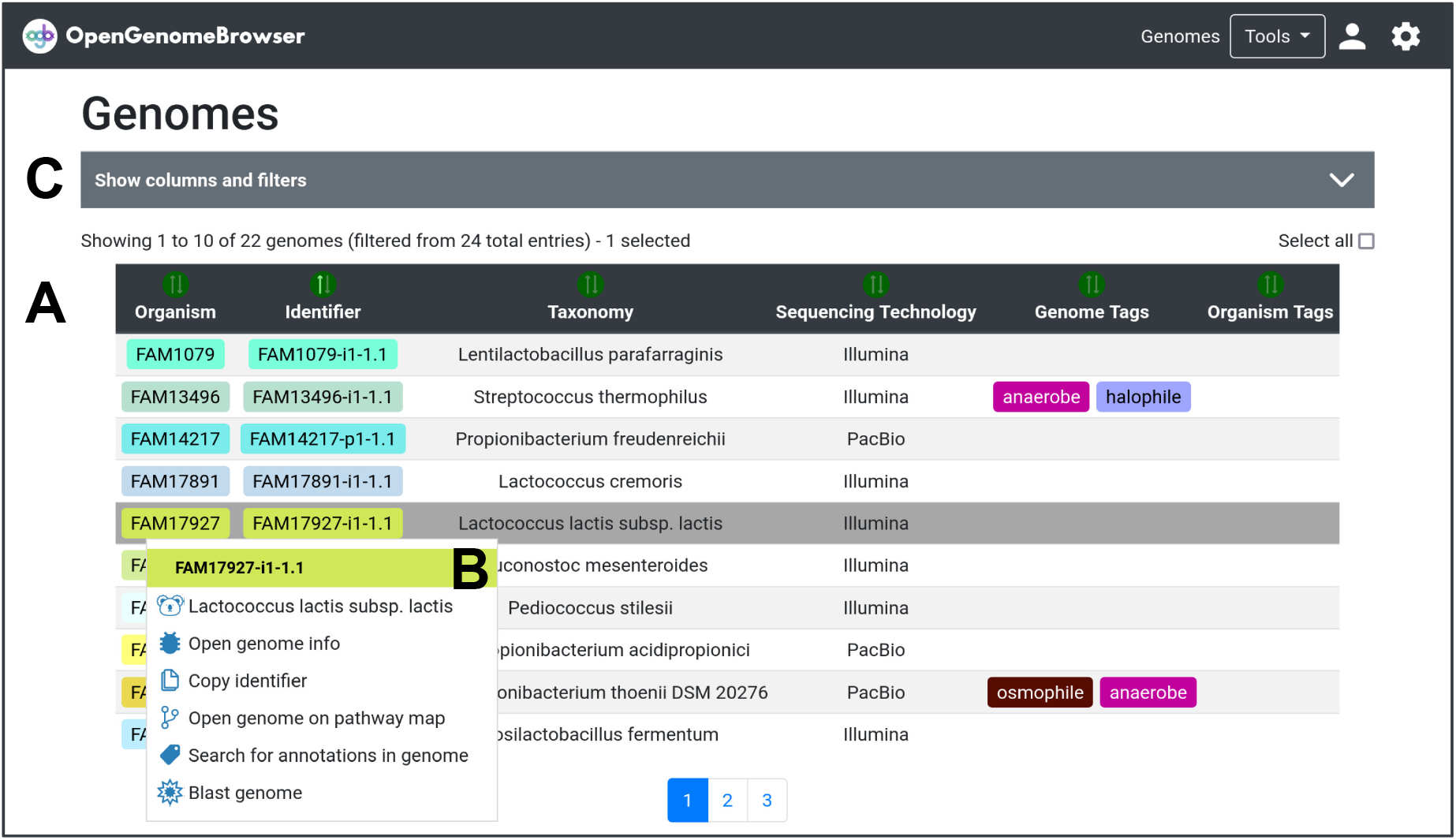
Genomes table. (**A**) Sortable and filterable table: Here, one genome is selected, but it is possible to select multiple. (**B**) Genome context menu: provides access to other features. (**C**) “Show columns and filters”: A click on this bar expands settings to add more metadata columns to the table and apply filters. (**⛭**) Settings sidebar: Download the table.

### Detail views

The *genome detail view* shows all available metadata of the respective genome and allows the user to download the associated data.

The *gene detail view* is designed to facilitate easy interpretation of the putative functions of genes. It shows all annotations, their descriptions, the nucleotide- and protein sequences, metadata from the GenBank file and an interactive gene locus visualization facilitated by DNA features viewer (19). If the gene is annotated with a gene ontology term that represents a subcellular location, this location will be highlighted on a SwissBioPics image (20).

Genomes in OpenGenomeBrowser can be labelled with tags, i.e., a short name (e.g., “*halophile*”) and a description (e.g., “*extremophiles that thrive in high salt concentrations*”). The *tag detail view* shows the description of the tag and the genomes that are associated with it. Tags are particularly useful to quickly select groups of genomes in many tools of OpenGenomeBrowser. For example, to select all genomes with the tag “*halophile*”, the syntax “*@tag:halophile*” can be used.

Similarly, the *TaxId detail view* shows all genomes that belong to the respective NCBI Taxonomy identifier (TaxId) (21), as well as the parent TaxId. Similar to tags, TaxIds can be used to select all genomes that belong to a certain TaxId, like this: “*@taxphylum:Firmicutes*”, or simply “*@tax:Firmicutes*”.

### Gene comparison

The *gene comparison view (*Figure 3) enables users to easily compute multiple sequence alignments and to compare gene loci side-by-side. Currently, Clustal Omega (22), MAFFT (23) and MUSCLE (24) are supported alignment algorithms. Alignments are visualized using MSAViewer (25) (Figure 3B). Furthermore, the genomic regions around the genes of interest can be analyzed using a customized implementation of DNA features viewer (19) (Figure 3C). Figure 3 shows an alignment of all genes on the demo server that contain the annotation *K01610* (phosphoenolpyruvate carboxykinase; from the pyruvate metabolism pathway). The gene loci comparison reveals that in all queried *Lacticaseibacilli*, the genes are located in syntenic regions, i.e., next to the same orthologous genes.

**Figure 3:**
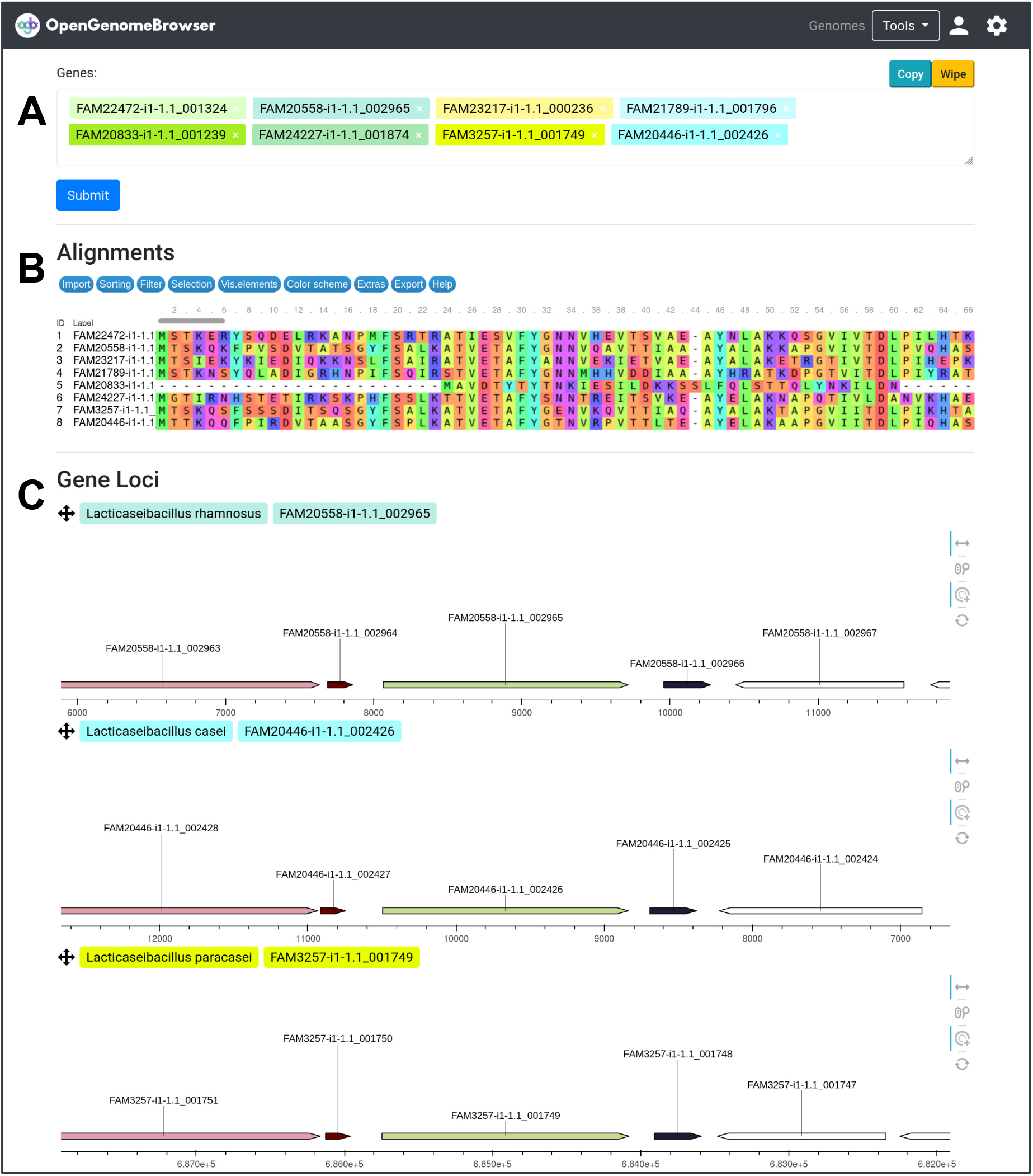
Gene comparison. (**A**) input mask for genes to be searched. (**B**) Output: alignments. Can be exported in aligned FASTA format. (**C**) Output: gene loci. Each subplot shows the genes around one of the queried genes, which are represented as colorful arrows. Orthologous genes have the same colors while genes without orthologs are white. The plots are interactive: pan, zoom, and click on genes. (**⛭**) Settings sidebar. For alignments: Choose between DNA and the protein sequence alignments, change the alignment method. For gene loci: Set the range around the gene’s locus, change the annotation category by which to color the genes.

### Annotation search

Despite conceptually and technically straightforward, searching for annotations in a set of genomes can be tedious or even impossible for non-programmers. In OpenGenomeBrowser, annotation search is quick and easy, thanks to the PostgreSQL backend that allows fast processing of annotation information. In the *annotation search view* (Figure 4), users can search for annotations in genomes, resulting in a *coverage matrix* (Figure 4C) with one column per genome and one row per annotation. The numbers in the cells show how many genes in the genome have the same annotation. Clicking on these cells shows the relevant genes (Figure 4D), while clicking on an annotation enables users to compare the corresponding genes (*gene comparison view)*.

**Figure 4:**
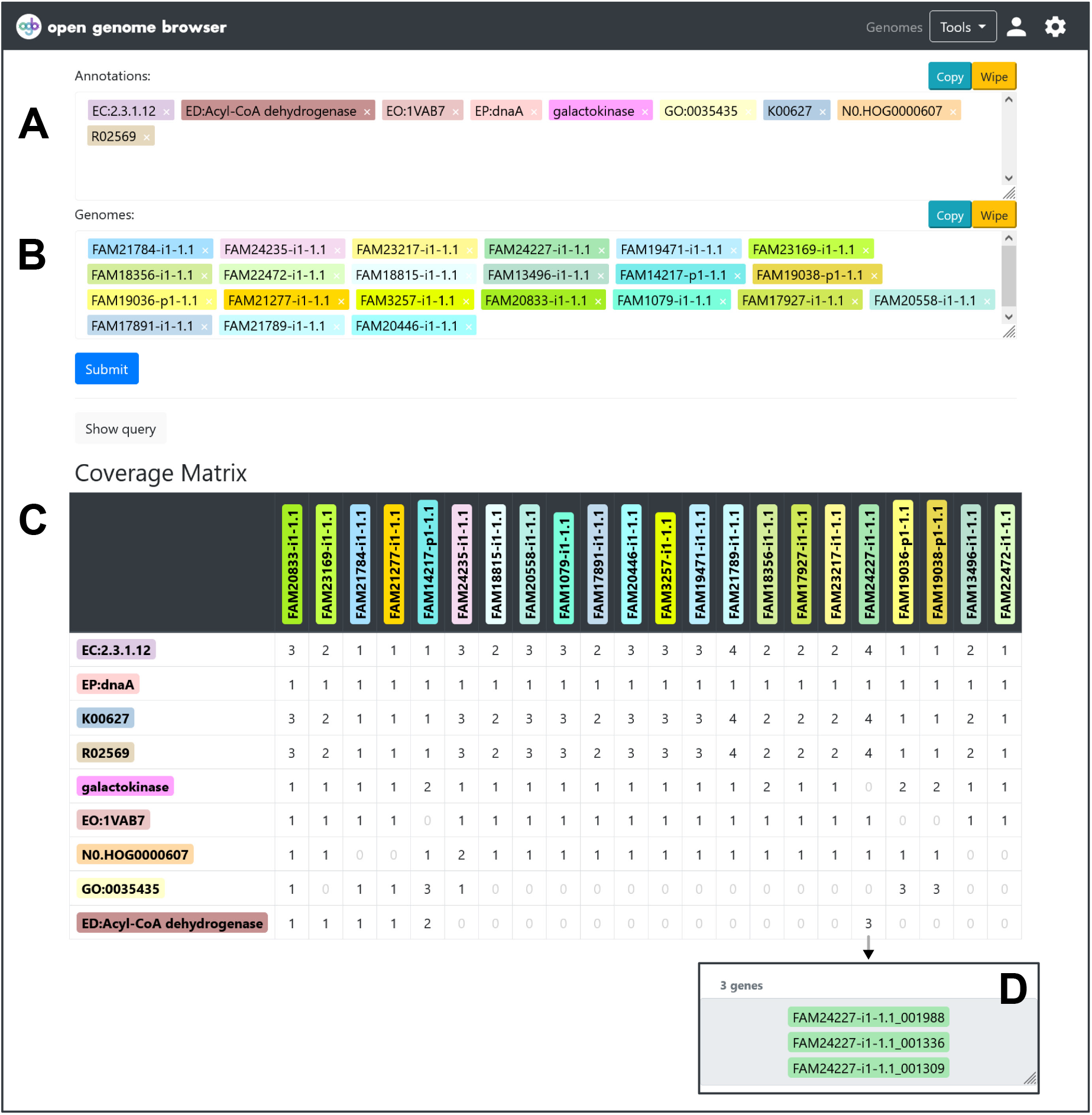
Annotation search. (**A**) Search for annotations. (**B**) Search for genomes. (**C**) *Coverage matrix* output: The numbers in the cells tell the number of genes in the genome that have the annotation. (**D**) Clicking on a cell reveals which genes. (**⛭**) Settings sidebar: Download the table.

### Pathways

Pathway maps, particularly the ones from the KEGG (26), are valuable tools to understand the metabolism of an organism. However, using them may be cumbersome. Commonly, biologists upload sequences to a service like BlastKOALA (27). This service is designed to process one organism at a time, and calculation times can last multiple hours. Because each genome must be submitted individually, it becomes cumbersome when multiple organisms must be processed. Furthermore, it is not trivial to visualize multiple genomes on a pathway map. In OpenGenomeBrowser, this process is straightforward, user-friendly (Figure 5A), and fast, as the annotations are pre-calculated and loaded into the database beforehand. Pathway maps are interactive, which allows the user to explore this information in great detail (Figure 5F). For example, to investigate the genes that are involved in a certain enzymatic step, one needs only to click on the enzyme box, then on an annotation of interest, and finally on “compare the genes” to be redirected to *gene comparison view*.

**Figure 5:**
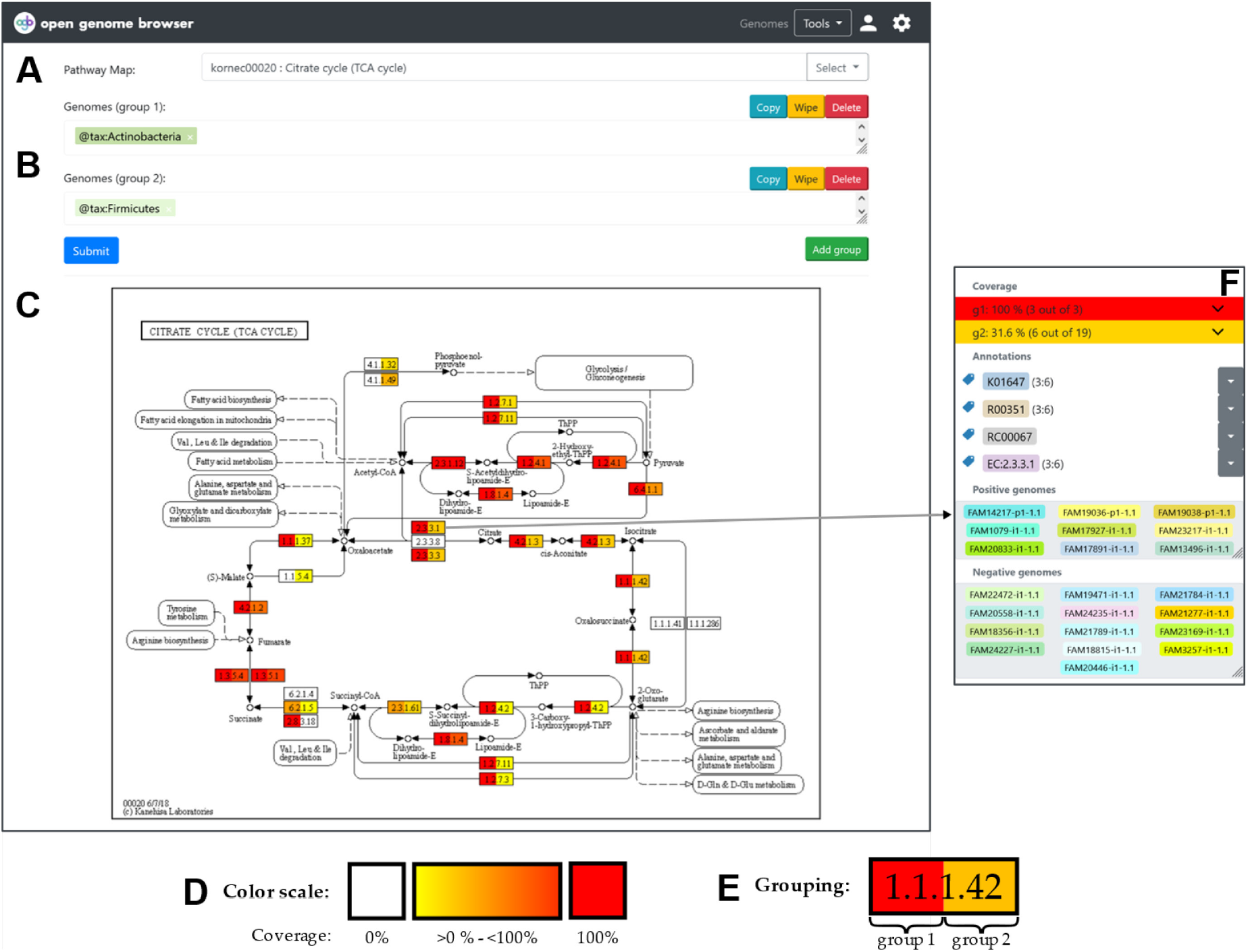
Pathway. Visualization of pathway coverage of one or multiple genomes. (**A**) Search for pathway maps. (**B**) Search for genomes: One or more groups of genomes can be added. In this example, the first group includes all genomes that belong to the taxonomic group *Actinobacteria*, the second group all *Firmicutes*. (**C**) Output: Coverage visualized on the KEGG citrate cycle pathway. (**D**) Color scale: The color of the reaction boxes indicates how many of the selected genomes cover the reaction (white: no genomes, yellow to red: one to all genomes). (**E**) Grouping: The left part of each box corresponds to the first group, the right part to the second. In this example, all *Actinobacteria* genomes (3 out of 3) in the database cover the entire citrate cycle, whereas not all *Firmicutes* (6 out of 19) do. (**F**) Context menu of a reaction box: It shows which annotations are behind the reaction and which genomes cover the reaction. (**⛭**) Settings sidebar: Change the colors, export the information in the plot as a table, download the pathway map in PNG or SVG format.

While OpenGenomeBrowser does not include KEGG maps for licensing reasons, users with appropriate rights can generate them using a separate program (28). The pathway maps do not necessarily have to be from KEGG. Pathway maps in a custom Scalable Vector Graphics (SVG) may be added to a designated folder in the folder structure (not shown in Figure 1.)

### BLAST

OpenGenomeBrowser allows users to perform a local alignment of protein and nucleotide sequences using BLAST (4). The results are visualized using the BlasterJS (29) library.

### Trees

OpenGenomeBrowser computes three kinds of phylogenetic trees. The fastest type of tree is based on the NCBI taxonomy ID which is registered in the metadata. It is helpful to get a quick taxonomic overview, but it entirely depends on the accuracy of the metadata.

The second type of tree is based on genome similarity. The assemblies of the selected genomes are compared to each other using GenDisCal-PaSiT6, a fast, hexanucleotide-frequency-based algorithm with similar accuracy as average nucleotide identity (ANI) based methods (30). This algorithm yields a similarity matrix from which a dendrogram is calculated with the unweighted pair group method with arithmetic mean (UPGMA) algorithm (31). We recommend this type of tree as a good compromise between speed and accuracy, specifically if many genomes are to be compared.

The third type of tree is based on the alignment of single-copy orthologous genes. This type of tree is calculated using the OrthoFinder (32) algorithm. Of all proposed tree type algorithms it is the most time- and computation-intensive and requires pre-computed all-vs-all DIAMOND (33) searches.

### Dot plot

Dot plot is a simple and established (34) method of comparing two genome assemblies. It allows the discovery of insertions, deletions, and duplications, especially in closely related genomes sequenced with long-read technologies. In OpenGenomeBrowser’s implementation of dot plot, the assemblies are aligned against each other using MUMmer (35) and visualized using the *Dot* library (36). The resulting plot (Figure 6) is interactive, i.e., the user can zoom in on regions of interest by drawing a rectangle with the mouse and clicking on a gene which then opens the context menu with detailed information.

**Figure 6:**
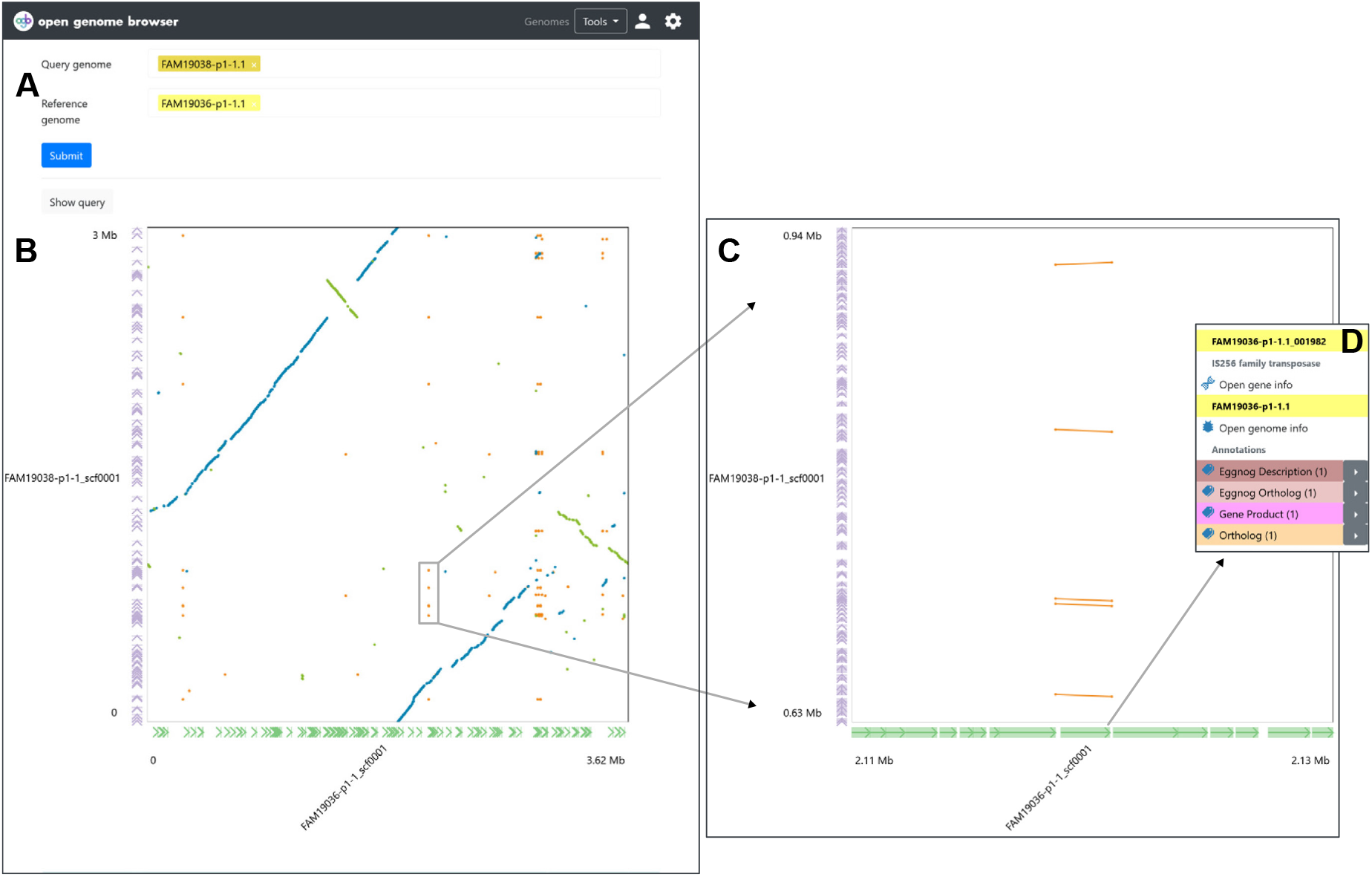
Dot plot view. (**A**) Search for query and reference genomes. (**B**) Output: dot plot. The reference genome is on the X-axis, the query genome on the Y-axis. The genes are shown at the edge of the plot in green and violet, respectively. By default, unique forward alignments are colored in blue, unique reverse alignments in green and repetitive alignments in orange. (**C**) Zooming in is achieved by drawing a rectangle with the mouse over a region with repeated elements. (**D**) A click on the suspicious gene reveals it to be a transposase. (**⛭**) Settings sidebar: Configure how the plot is rendered.

### Gene trait matching

The *gene trait matching view* enables users to find annotations that correlate with a (binary) phenotypic trait. The input must consist of two non-intersecting sets of organisms that differ in a trait. OpenGenomeBrowser applies a Fisher’s exact test for each orthologous gene and corrects for multiple testing (alpha = 10%) using the Benjamini-Hochberg method (37,38). The multiple testing parameters can be adjusted in the settings sidebar. The test can be used on orthogenes as well as any other type of annotation, such as KEGG-gene annotation. The gene candidates that may be causing the trait can easily be further analyzed, for example by using the *compare genes view*.

### Flower plot

The *flower plot view* provides the users with a simple overview of the shared genomic content of multiple genomes. The genomes are displayed as petals of a flower. Each petal indicates the number of annotations that are unique to this genome and the number of genes that are shared by some but not all others. The number of genes shared by all genomes is indicated in the center of the flower. (The code is also available as a standalone Python package. (39))

### Downloader

The *downloader view* facilitates the convenient download of multiple raw data files, for example all protein FASTA files for a set of organisms.

### Admin panel

OpenGenomeBrowser has a powerful user authentication system and admin interface, inherited from the Django framework. Instances of OpenGenomeBrowser can be configured to require a login or to allow basic access to anonymous users. Users can be given specific permissions, for example to create other user accounts, to edit metadata of organisms, genomes, and tags, and even to upload new genomes through the browser.

## Conclusions

OpenGenomeBrowser is, to our knowledge, the first comparative genome browser that is not tied to a specific dataset. It automates commonly used bioinformatics workflows, enabling convenient and fast data exploration, particularly for non-bioinformaticians, in an intuitive and user-friendly way.

The software has minimal hardware requirements and is easy to install, host, and update. OpenGenomeBrowser’s folder structure enforces systematic yet flexible storage of genomic data, including associated metadata. This folder structure (i) enables automation of analyses, (ii) guides users to maintain their data in a coherent and structured way, and (iii) provides version tracking, a precondition for reproducible research.

OpenGenomeBrowser is flexible and scalable. It can run on a local machine or on a public server, access may be open for anyone or restricted to authenticated users. Annotation types can be customized, and ortholog-based features are optional. While the demo server only holds 24 genomes, the performance scales and is still outstanding even when hosting over 1’400 microbial genomes (40).

We believe that our software will be useful to a large community since sequencing microbial and other genomes has become a commodity. Therefore, researchers performing new sequencing projects can directly benefit from OpenGenomeBrowser by saving development costs, making their data potentially FAIR, and adapting the browser for their purposes. It could also replace older, custom-made platforms which may be outdated and more difficult to maintain. Because our software is open-source, adaptations of OpenGenomeBrowser and new features will be available for the whole community under the same conditions. The open-source model also allows problems to be identified and quickly fixed by the community, making OpenGenomeBrowser a sustainable platform.

## Methods

### Software requirements

The only prerequisites to run an OpenGenomeBrowser server are a 64-bit Linux system with Docker and Docker Compose. OpenGenomeBrowser’s numerous software dependencies are contained in the Docker image. We provide installation instructions and detailed documentation at opengenomebrowser.github.io.

### Hardware requirements

OpenGenomeBrowser is not resource intensive. An instance containing over 1,400 bacterial genomes runs on a computer with 8 CPU-cores (2.4 GHz) and 20 GB of RAM. The Docker container is about 3 GB in size and the Postgres database takes 21 GB of storage (SSD recommended).

## DECLARATIONS

### ETHICS APPROVAL AND CONSENT TO PARTICIPATE

Not applicable.

## CONSENT FOR PUBLICATION

Not applicable.

## AVAILABILITY OF DATA AND MATERIALS

Project name: OpenGenomeBrowser

Project home page: https://opengenomebrowser.github.io/

Archived version: https://github.com/opengenomebrowser/opengenomebrowser

Operating system(s): Linux

Programming language: Python

Other requirements: Docker

Any restrictions to use by non-academics: GPL-3

## COMPETING INTERESTS

The authors declare that they have no competing interests.

## FUNDING

This research was funded by Gebert Rüf Stiftung within the program “Microbials”, grant number GRS-070/17 and the Canton of Bern to RB.

## AUTHORS’ CONTRIBUTIONS

TR, SO and RB conceived the project. TR programmed the software. SO, RB and NS contributed conceptually and with feedback to the software. TR and RB wrote the manuscript. All authors edited, read, and approved the final manuscript.

## ACKNOWLEDGEMENTS

We are grateful to Darja Studer for designing the logo, to Lars Vögtlin for his advice on containerization, to Linda Studer for her advice on the manuscript, and to Kimberly Gilbert for proofreading the article. We thank Emmanuelle Arias-Roth, Remo Schmidt, Cornelia Bär, Ueli von Ah und Guy Vergères (Agroscope) for their support and feedback for this project.

## Notes

https://opengenomebrowser.github.io/

https://opengenomebrowser.bioinformatics.unibe.ch/

### Competing Interest Statement

The authors have declared no competing interest.

